# Somatic Cell Nuclear Transfer Embryos Show Massive Dysregulation of Genes Involved in Transcription Pathways

**DOI:** 10.1101/2020.12.06.413393

**Authors:** Chunshen Long, Hanshuang Li, Xinru Li, Yongchun Zuo

**Author notes:** These authors contributed equally: Chunshen Long and Hanshuang Li.

## Abstract

Transcription is the most fundamental molecular event that occurs with zygotic genome activation (ZGA) during embryo development. However, the potential association between transcription pathways and low cloning efficiency of nuclear transfer (NT) embryos remains elusive. Here, we integrated a series of RNA-seq data on NT embryos to deciphering the molecular barriers of NT embryo development. Comparative transcriptome analysis indicated that incomplete activation of transcription pathways functions as a barrier for NT embryos. Then, the gene regulatory network (GRN) identified that crucial factors responsible for transcription play a coordinated role in epigenome erasure and pluripotency regulation during normal embryo development. But in NT embryos, massive genes involved in transcription pathways were varying degrees of inhibition. Our study therefore provides new insights into understanding the barriers to NT embryo reprogramming.

## Introduction

Somatic cell nuclear transfer (SCNT) technology can reprogram terminally differentiated cell nuclei into a totipotent state to realize the cloning of animals. SCNT has great prospects in therapeutic cloning, animal breeding and endangered species protection ^[1-4]^. At present, there are still many technical obstacles in somatic cell nuclear transfer, which make SCNT embryos have low cloning efficiency, extra-embryonic tissues and some abnormal phenomena after the birth of cloned animals ^[5, 6]^. In mouse, 70% of SCNT embryos are arrested at early cleavage stages, especially from the 1C to the 2C stage ^[7, 8]^. And a recent analysis indicates that inhibition of minor ZGA impairs the Pol II pre-configuration and embryonic development in mouse embryos^[9]^. Thus, whether transcription-related pathways also play roles in SCNT embryos needs to be further investigated. In accordance with our previous study, we observed that the transcripts related to transcription, such as TFIID subunits, RNA polymerase and mediators as the main trigger genes, which are not fully activated in inter-SCNT embryos ^[10]^.

## Result and Discussion

Transcription is one of the most fundamental cellular events and the first occurrence of this process is accompanied by the zygotic genome activation (ZGA). However, the potential influence of transcription-related pathways on embryos development remains elusive. To address this, we collected three pathways related to transcription-related pathways from the KEGG database. These three KEGG pathways are Basal transcription factors (TFs), RNA polymerase and Spliceosome, involving 44, 31, and 132 factors respectively. In order to further explore the developmental defects of SCNT embryos from the perspective of transcription-related pathways activation, we compared the activation differences of Basal TFs, RNA polymerase and Spliceosome between *in vivo* fertilized embryos (WT) and SCNT embryos (Fig. 1a). Unexpectedly, the defective activation of these three pathways was observed in SCNT embryos compared to *in vivo* fertilized embryos, showing that incomplete activation of transcription pathways may be the cause of development arrest of SCNT embryos. Consistently, both in nuclear transfer 2 and 4-cell arrest (NTA2 and NTA4) embryos, the expression levels of genes involved in Basal TFs, RNA polymerase and Spliceosome are significantly lower than corresponding development stages of *in vivo* fertilized embryos (WT2 and WT4) (Fig. 1b).

**Fig. 1.**
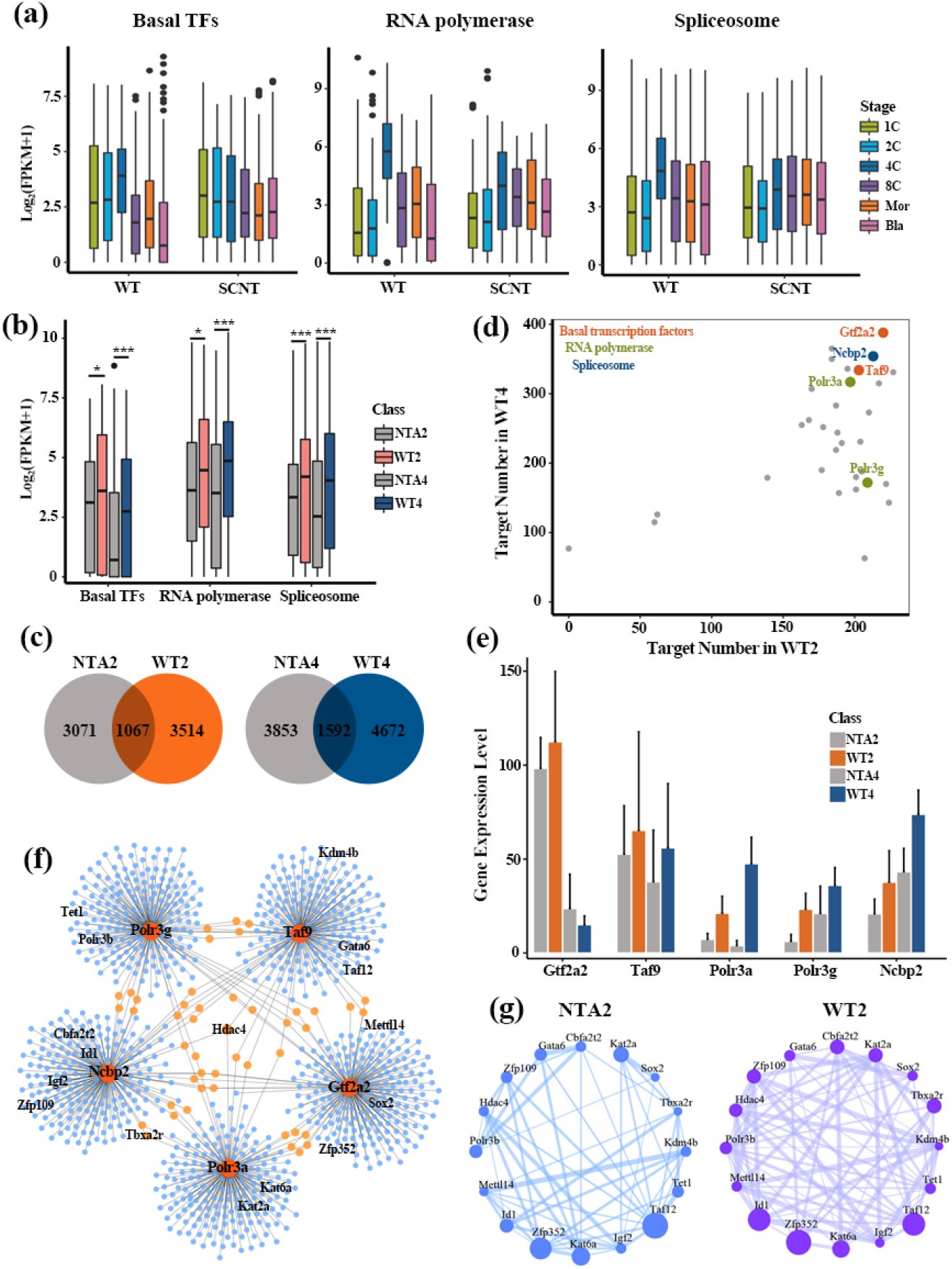
Incomplete activation of transcription-related pathways in SCNT embryos. **(a)** Abnormal activation of three key pathways (Basal TFs, RNA polymerase and Spliceosome) related to basal transcription event was observed in SCNT embryos. **(b)** The boxplot shows the differential activation of three pathway between in NT arrest and normal embryos. Differences are statistically significant. (*) P-value < 0.05; (**) P-value < 0.01; (***) P-value < 0.001, t-test. NTA2, NT 2-cell arrest embryos; WT2, *in vitro* fertilization 2-cell embryos; NTA4, NT 4-cell arrest embryos; WT4, *in vitro* fertilization 4-cell embryos. **(c)** Overlap between targets genes regulated by three pathway related TFs in NT arrest and normal development embryos. **(d)** The selection of candidate TFs (marked in figure) involved in three pathways, the five TFs having top targets both in 2 and 4 cell embryos were screened. **(e)** The differential expression patterns of five representative genes between in NT arrest and normal embryos. The expression patterns of these genes are represented as the average plus standard deviation (SD) of biological replicates (Mean+SD). **(f)** The transcription networks of the five TFs regulating the normal development of 2cell embryos. **(g)** The expression networks for the target genes (marked in Figure 1E) of five hub factors in NTA2 and WT2 embryos, respectively. The dot size represents the gene expression level and the connection line indicates Pearson correlation.

The above data have indicated that the existence of aberrant transcription process was significantly linked to embryos development (Fig. 1a and 1b). Thus, we wondered whether the dynamic activation of transcription-related pathways could provide clues for identifying key factors mediating the normal development of SCNT embryos. To this end, we focused on the differences of regulatory network between normal development embryos and development arrest embryos. Based on pySCENIC (python package), we constructed the gene regulatory network (GRN) involved in transcription-related factors in 2 and 4-cell embryos with different development fates. Compared with NT 2-cell arrest (NTA2) embryos, the more target genes have been observed in regulatory network of *in vitro* fertilization 2-cell (WT2) embryos, with consistent results in WT4 embryos (Fig. 1c). These findings implied that the successful activation of gene regulatory networks involved in transcription contributes to the normal development of SCNT embryos.

Given that the correct activation of the transcriptional regulatory network is essential for the normal development of SCNT embryos, we next sought to identify pioneer factors with key roles from genes of transcription pathways. And the top 1 or 2 core factors with the largest targets number in WT embryos were screened, including Gtf2a2, Taf9 (Basal TFs), RNA Polr3a, Polr3g (polymerase) and Ncbp1 (Spliceosome) (Fig. 1d). Compared with normal developmental embryos, massive dysregulation of genes involved in transcription pathways was observed in NT arrest embryos (except for Gtf2a2 in NTA4 embryos) (Fig. 1e).

To understand why the repression of these genes associate with embryos arrest, we focus our research on 2-cell embryos and constructed a regulatory network with the above five factors as hub genes in WT2 embryos (Fig. 1f). Interestingly, these regulatory factors are specifically present in WT2 embryos, and no corresponding regulatory relationships have been observed in NTA2 embryos. Among them, Hdac4, as a histone deacetylase, is co-regulated by Taf9, Polr3a and Ncbp1, suggesting the crucial epigenetic control defects in SCNT embryo development. Histone H3K9me3 demethylase Kdm4b was targeted by Taf9, which is consistent with previous studies that Kdm4b functions as a barrier for NTA2 embryos ^[8]^. Moreover, DNA demethylase Tet1 ^[8, 11]^, histone acetylase Kat2a and Kat6a were also observed in this GRN, both of which are highly expressed in WT2 embryos (Fig. 1g). Additionally, we also found that some pluripotency factors, such as Sox2, Taxa2r, Cbfa2t2, Id1, Zfp109 and Zfp352, have higher expression levels in the WT2 embryos compared to NTA2 embryos (Fig. 1g). This result shows that the maintenance of pluripotency is essential for normal embryo development. Notably, Gata6 and Igf2 may serve as candidates for potential activation of defects in SCNT embryos and promote SCNT efficiency.

## Conclusion

Here, we identified massive dysregulation of genes related to transcription pathway in NT embryo. And GRN further indicated that these genes play a coordinated role in erasing the epigenetic barrier, enhancing pluripotency regulation to facilitate embryos development. Overall, our study provides new clues for improving the efficiency of SCNT reprogramming from the perspective of transcription-related pathways activation.

## Materials and Methods

### Dataset Collection

The single-cell RNA sequencing (RNA-seq) data of mouse pre-implantation embryos development was downloaded from Gene Expression Omnibus (GEO) database under accession number GEO: GSE113164^[12]^. There are two embryonic types, including somatic cell nuclear transfer (SCNT) embryos and *in vitro* fertilization (WT) embryos. Both SCNT and WT samples include Zygote, 2-cell, 4-cell, 8-cell, Morula, Blastocyst, and each stage has three replicates. Moreover, another single-cell RNA-seq data (GSE70605) ^[8]^ was also reanalyzed in this study, which includes two embryos types SCNT embryos and *in vitro* fertilization embryos. And these embryos can be divided into nuclear transfer (NT) 2cell arrest embryos (NTA2), NT 4cell arrest embryos (NTA4), *in vitro* fertilization 2cell embryos (WT2) and *in vitro* fertilization 4cell embryos (WT4).

### RNA-seq Data Processing

For RNA-seq data processing, all RNA-seq data was controlled by FastQC software (http://www.bioinformatics.babraham.ac.uk/projects/fastqc/) and raw reads were trimmed based on Trimmomatic (version 0.38) ^[13]^ to remove low quality samples. Next, filtered reads were mapped to the mouse mm9 genome with HISAT2 (version 2.1.0) [14] aligner with default parameters. Then, read counts of each gene were calculated using HTseq (version 0.11.0) ^[14]^. Transcriptome assembly was performed using Stringtie (version 1.3.3) ^[14, 15]^ and Ballgown (R package), and expression levels of each gene were quantified with normalized FPKM (fragments per kilobase of exon model per million mapped reads) ^[16]^.

### Functional Pathways Activation Analysis

Basal TFs, RNA polymerase and Spliceosome related to basic transcription were obtained from Kyoto Encyclopedia of Genes and Genomes (KEGG) database (https://www.kegg.jp/kegg/pathway.html) ^[17]^. And the activation of these three KEGG pathways during pre-implantation embryos development was visualized by boxplot.

### Single-cell Regulatory Network Inference

The workflow of pySCENIC ^[18]^ (https://pypi.org/project/pyscenic/0.6.6/#tutorial) was used to study the gene regulatory network involved in basic transcription-related factors during embryonic development. In pySCENIC workflow, RcisTarget ^[19]^ package determine TFs and their target genes (i.e. targetomes) based on the correlation of gene expression across cells, and GRNBoost ^[20]^ identify whether the target genes have the corresponding motifs of TF to refine targetomes. Finally, active targetomes are recognized in every single cell. The regulatory network centered on basic transcription-related factors was screened out and visualized by Cytoscape ^[21]^.

### Data Visualization

In this study, R/Bioconductor (http://www.bioconductor.org) software packages were mainly used to data visualization. For example, the Venn plot was produced by using R packet VennDiagram. And the barplot, boxplot and scatter plot were generated with the R packet ggplot2 (http://ggplot2.org/).

## Conflict of interest

The authors declare that they have no conflict of interest.

## Acknowledgments

We thank the financial support from the National Nature Scientific Foundation of China (grants 62061034, 61702290 and 61861036); Program for Young Talents of Science and Technology in Universities of Inner Mongolia Autonomous Region (grant NJYT-18-B01); and the Fund for Excellent Young Scholars of Inner Mongolia (grant 2017JQ04).

## Author Contribution

Yongchun Zuo conceived and designed the study. Chunshen Long and Hanshuang Li conducted the experiments. Xinru Li contributed to partial characterizations. Yongchun Zuo and Chunshen Long wrote the paper. All authors read and approved the manuscript.

## Notes

### Competing Interest Statement

The authors have declared no competing interest.

### Summary of Updates

Figure 1 revised

